# Decade-long remissions of leukemia sustained by the persistence of activated CD4^+^ CAR T-cells

**DOI:** 10.1101/2021.05.07.443194

**Authors:** J. Joseph Melenhorst, Gregory M. Chen, Meng Wang, David L. Porter, Peng Gao, Shovik Bandyopadhyay, Iulian Pruteanu-Malinici, Christopher L. Nobles, Sayantan Maji, Noelle V. Frey, Saar I. Gill, Lifeng Tian, Irina Kulikovskaya, Minnal Gupta, Megan M. Davis, Joseph A. Fraietta, Jennifer L. Brogdon, Regina M. Young, David E. Ambrose, Anne Chew, Bruce L. Levine, Donald L. Siegel, Cécile Alanio, E. John Wherry, Frederic D. Bushman, Simon F. Lacey, Kai Tan, Carl H. June

**Affiliations:** Center for Cellular Immunotherapies, Perelman School of Medicine, University of Pennsylvania, Philadelphia, PA, USA; Abramson Cancer Center, Perelman School of Medicine, University of Pennsylvania, Philadelphia, PA, USA; Department of Pathology and Laboratory Medicine, Perelman School of Medicine, University of Pennsylvania, Philadelphia, PA, USA; Institute for Immunology, Perelman School of Medicine, University of Pennsylvania, Philadelphia, PA, USA; Graduate Group in Genomics and Computational Biology, University of Pennsylvania, Philadelphia, PA, USA; Graduate Group in Cell & Molecular Biology, University of Pennsylvania, Philadelphia, PA, USA; Center for Childhood Cancer Research, The Children’s Hospital of Philadelphia, Philadelphia, PA, USA; Department of Pediatrics, Perelman School of Medicine, University of Pennsylvania, Philadelphia, USA; Novartis Institute for Biomedical Research, Cambridge, MA, USA; Department of Microbiology, Perelman School of Medicine, University of Pennsylvania, Philadelphia, PA, USA; Department of Transfusion Medicine, Perelman School of Medicine, University of Pennsylvania, Philadelphia, PA, USA; Department of Systems Pharmacology and Translational Therapeutics, Perelman School of Medicine, University of Pennsylvania, Philadelphia, PA, USA; Parker Institute for Cancer Immunotherapy, University of Pennsylvania, Philadelphia, USA

## Abstract

The adoptive transfer of T lymphocytes reprogrammed to target tumor cells has demonstrated significant potential in various malignancies. However, little is known about the long-term potential and the clonal stability of the infused cells. Here, we studied the longest persisting CD19-redirected chimeric antigen receptor (CAR) T cells to date in two chronic lymphocytic leukemia (CLL) patients who achieved a complete remission in 2010. CAR T-cells were still detectable up to 10+ years post-infusion, with sustained remission in both patients. Surprisingly, a prominent, highly activated CD4^+^ population developed in both patients during the years post-infusion, dominating the CAR T-cell population at the late time points. This transition was reflected in the stabilization of the clonal make-up of CAR T-cells with a repertoire dominated by few clones. Single cell multi-omics profiling via Cellular Indexing of Transcriptomes and Epitopes by Sequencing (CITE-Seq) with TCR sequencing of CAR T-cells obtained 9.3 years post-infusion demonstrated that these long-persisting CD4^+^ CAR T-cells exhibited cytotoxic characteristics along with strong evidence of ongoing functional activation and proliferation. Our data provide novel insight into the CAR T-cell characteristics associated with long-term remission in leukemia.

## Main Text

By reprogramming patient T lymphocytes with a chimeric antigen receptor (CAR) specific for CD19, we^1–4^ and others^5–7^ demonstrated that durable remissions are attainable in relapsed, refractory B cell leukemias and lymphomas. The first two CLL patients were infused in the summer of 2010 with anti-CD19 CAR T (CTL019) cells, and responded with complete remissions and persistence of the infused CAR T-cells^1^. Persistence of CAR T-cells in acute lymphoid leukemia (ALL) and CLL is a key characteristic of durable clinical responses^2,8–10^, yet the characteristics of long-term persisting CAR T-cells have not been extensively studied. The particular persistence of anti-CD19 CAR T-cells in these first two complete-responding CLL patients allowed us to interrogate molecular and functional attributes of highly effective anti-CLL T cells. Here, we report that these two patients have remained in remission at last follow-up >10 years post-infusion, and we have mapped the fate of CTL019 cells using bulk and single-cell multi-omic approaches.

## Results

### Sustained CLL remissions with CAR T-cells detected 10 years post-infusion

Two patients were infused with autologous CAR T-cells as part of a phase I clinical trial in the summer of 2010^1–4^. Peak CTL019 expansion in patient 1 occurred at day 3, whereas this was delayed to day 31 in patient 2, a possible reflection of the almost 78-fold lower infusion dose. Here, we report that these patients have remained in remission >10 years post-infusion. CAR T-cells were detectable by flow cytometry across time points (**Fig. 1a-c**); at the time of the most recent research phlebotomy, which was at 10 years post-infusion for patient 1 and 9 years for patient 2, CTL019 cells remained detectable and represented 0.8% and 0.1% of all T-cells respectively (**Fig. 1b-c**). Quantitative PCR confirmed the presence of CTL019 in both patients at all time points. Using flow cytometry, CD19^+^ B-lymphocytes and CLL cells have been undetectable or highly suppressed (<<1% of cells) beyond 3 years post-infusion. The absence of leukemia cells was confirmed with immunoglobulin heavy (IgH) chain repertoire analysis: the leukemic clone has remained undetectable since 6 months after CTL019 administration. Further, productively rearranged IgH sequences have been at background levels since 12 and 6 months after anti-CD19 CAR T-cell treatment in patients 1 and 2, respectively (**Extended Data Table 1**), confirming the B cell aplasia findings by flow cytometry.

**Fig. 1.**
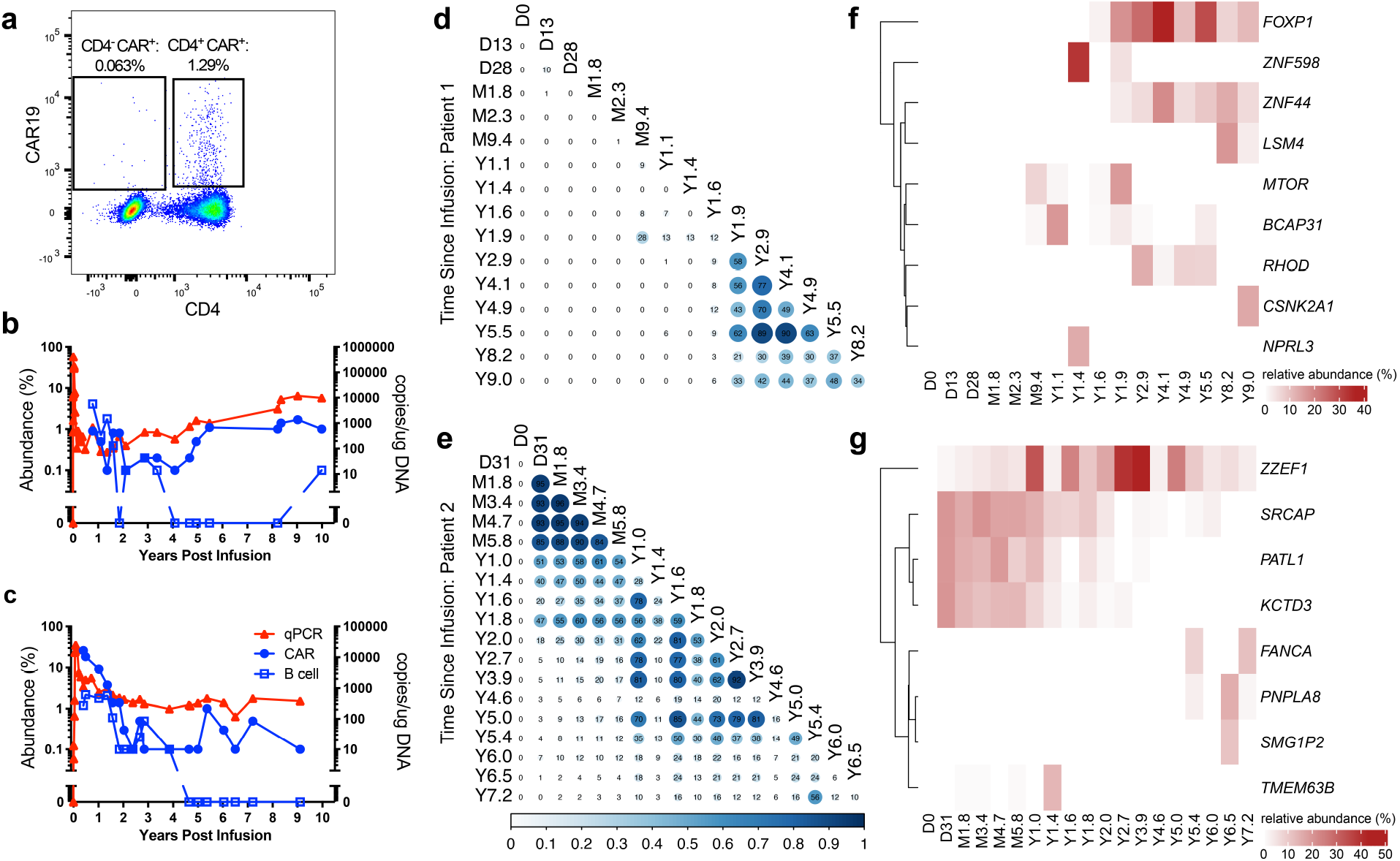
Molecular tracking of effectors and targets in long-term responders to anti-CD19 CAR T-cell therapy for CLL. **a**, Representative flow cytometry data displaying persistence of mostly CD4^+^ CTL019 cells in patient 1, eight years after infusion. **b-c**, Kinetics of CAR T-cell expansion and persistence (red triangles, by qPCR with vector-specific primers; blue circles, by flow cytometry using an anti-CAR antibody) and response of B cells (blue squares, by anti-CD19 flow cytometry) to anti-CD19 CAR T-cell therapy in patient 1 (top) and 2 (bottom). **d-e**, Clonal evolution of CAR T-cells based on lentiviral vector integration site analysis. Pairwise Morisita’s overlap index (shown as two decimal points in each circle) was computed between all timepoints (in days post-infusion) for patient 1 (**d**) and 2 (**e**). Integration sites with abundance > 10% in at least one time point were tracked over time for patient 1 (**f**) and 2 (**g**). The integration sites were labeled using the nearest genes of integration.

### Molecular fate mapping reveals clonal stabilization of the CAR T-cell repertoire

To understand the clonal nature and evolution of the CAR T-cell expansions, we examined the T cell receptor (TCR)-β chain repertoire of sorted CAR T-cells via deep sequencing (TCR-Seq) and lentiviral vector integration site (LVIS) analysis^11^ in both patients. TCR-Seq revealed a clonal shift in patient 1 that occurred between the month 2.3 and year 1.4 time points; in contrast to the clonal composition of patient 2, which was overall more stable with a gradual shift over the first two years (**Extended Data Fig. 1a-d**). TCR-Seq was limited to the early post-infusion time points due to input cell requirements, so we turned to LVIS analysis^11,12^ to assess both clonal architecture, dynamics, and CAR integration sites across time points up to 9.0 and 7.2 years post-infusion in patients 1 and 2 respectively. We identified 7,930 and 3,406 total unique sites for patients 1 and 2; 3,378 and 1,216 were identified in the infusion products of patient 1 and 2, respectively. Most insertions were present within the gene body, and intronic insertions were favored over other gene elements (**Extended Data Fig. 2a-c**). Consistent with the TCR-Seq analysis, our LVIS data revealed little if any CAR T-cell clonal stability in the first 1.6 years in patient 1. From year 1.9 onward, the LVIS repertoire in this patient stabilized and remained that way until the last follow-up (**Fig. 1d**). Patient 2 had episodes of repertoire stability from day 31 to approximately year 1, as well as from year 1 to 5 (**Fig. 1e**).

Patient 1 showed emergence of clones with CAR insertion sites into *MTOR*, *BCAP31*, *ZNF598*, and *NPRL3* from the month 9.4 to year 1.9 time points, mostly at single time points. CAR insertions into or near *FOXP1*, *ZNF44*, and *RHOD* emerged in the year 1.6-1.9 time points and were stable over many years; and CAR insertion sites in *LSM4* and *CSNK2A* emerged in the year 8.2 and 9.0 time points (**Fig. 1f**). Patient 2 exhibited multiple CAR T-cell clones at >10% abundance early after infusion, such as *SRCAP*, *PATL1*, and *KCTD3* which were most prominent in the first two years, and *ZZEF1* which was sustained for >7 years after infusion. Other insertions acquired prominence later after infusion, including *FANCA*, *PNPLA8*, *SMG1P2*, and *TMEM63B* (**Fig. 1g**). Our data therefore show that the long-term persisting CAR T-cell repertoire in both patients stabilized over time and was characterized by the pauci-clonal dominance with non-overlapping insertion sites.

### CyTOF analysis reveals distinct CAR T-cell phenotypes associated with initial response and long-term remission phases

We developed a 40-antibody panel to deeply interrogate the phenotype of CAR T-cells at multiple time points using Cytometry by Time-of-Flight (CyTOF). We stringently gated CD3^+^ CAR^+^ T cells (**Extended Data Fig. 3a**), recovering over 45,000 CD3^+^ CAR^+^ cells across all time points, represented by at least 100 cells from each of five time points per patient (**Fig. 2a-b, Extended Data Fig. 3b**). In patient 1, CD8^+^ cells constituted 29.3% of CAR T-cells at month 1.8 and diminished in proportion at subsequent time points, with CD4^+^ cells constituting 97.5% of CAR T-cells at year 1.4 and over 99.6% of CAR T-cells from year 3.4 to the latest time point at 9.3 years post-infusion (**Fig. 2a-c**). Patient 2, who had more delayed CAR T-cell expansion characteristics, exhibited an overall similar trend, with prominent CD8^+^ cells in the initial time points, subsequently diminishing to such an extent that CD4^+^ cells constituted 97.6% of CAR T-cells by 7.2 years post-infusion. We additionally observed a population of CD4^-^CD8^-^ double-negative CAR T-cells that was most prominent in patient 2, constituting 33.4% and 46.5% of CAR T-cells at month 2.5 and year 1.6 respectively and diminishing to 12.9%, 8.2%, and 0.5% at years 2.8, 4.7, and 7.2 respectively (**Fig. 2b-c**). This double-negative population expressed CD3, CAR, and CD45 at similar levels as the CD4^+^ and CD8^+^ T cells, and had a distinct cellular phenotype, expressing cytotoxic markers GZMB, 2B4, CD57, CD85j, T-bet, and PD-1; as well as Helios, which notably differentiated this population from otherwise similar cytotoxic CD8^+^ T cells (**Fig. 2b; Extended Data Fig. 3b**). Given this distinct marker profile, we labeled this subset as double-negative Helios^hi^ CAR T-cells. This population was detected at low levels (3.2% of cells) at the month 1.8 time point in patient 1, and was absent or rare (< 1% of cells) at and beyond the year 1.4 time point (**Fig. 2c**).

**Fig. 2.**
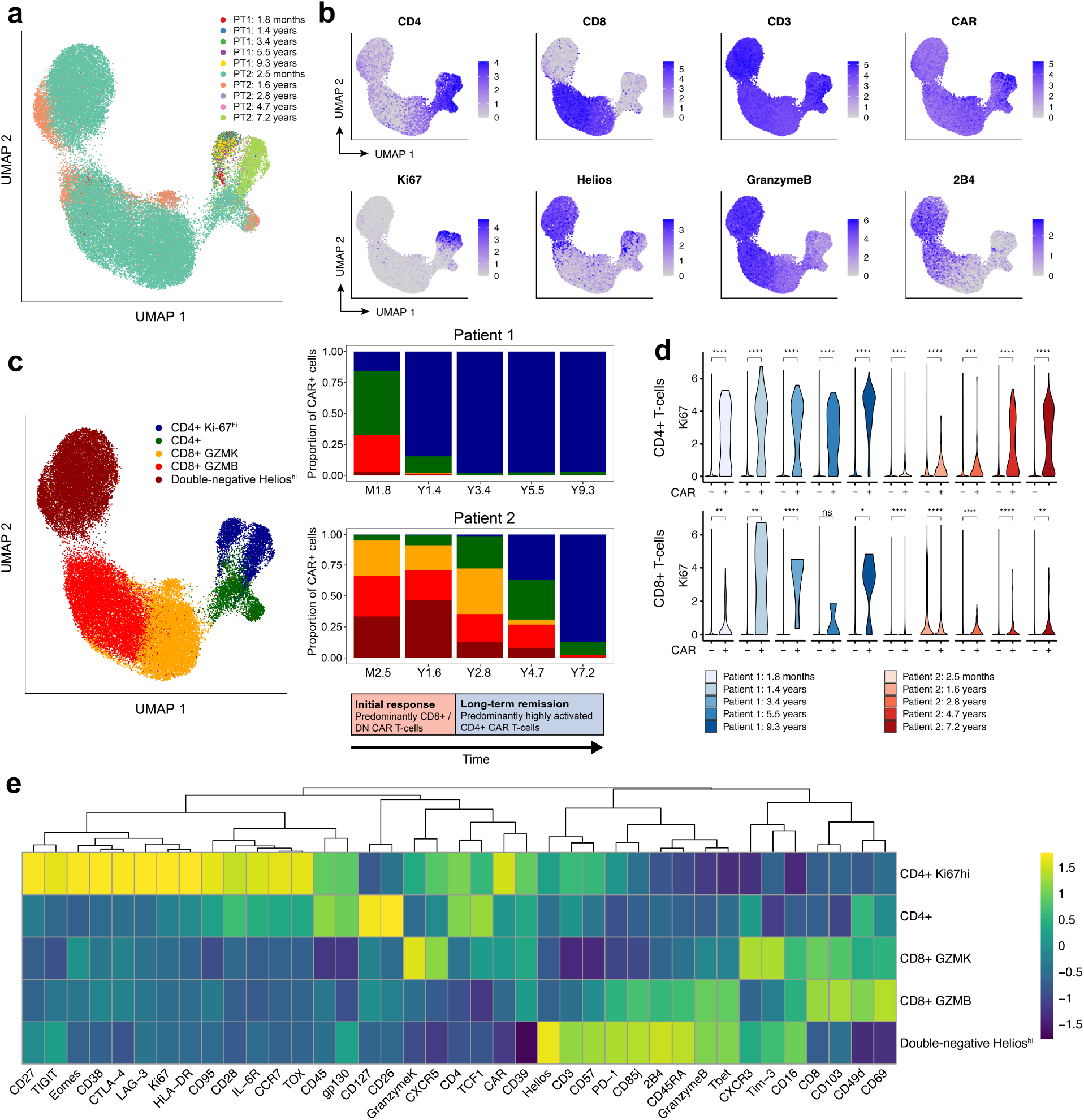
Analysis of CD3^+^ CAR^+^ T cells using CyTOF across multiple time points. **a,** UMAP of CD3^+^ CAR^+^ gated cells from CyTOF data generated from samples at five time points for each of patient 1 (PT1) and patient 2 (PT2). Each color represents cells from one patient time point post-infusion. **b**, Protein expression of selected CyTOF markers, revealing prominent Ki-67^hi^ population of CD4^+^ CAR T-cells, as well as a CD4^-^ CD8^-^ double-negative population expressing Helios, Granzyme B, and 2B4. **c**, UMAP grouped by five major clusters of CAR T-cells: a CD4^+^ Ki-67^hi^ population, a CD4^+^ population without this Ki-67^hi^ phenotype, a CD8^+^ population highly expressing Granzyme K, a CD8^+^ population highly expressing Granzyme B, and a double-negative population expressing Helios. The adjacent stacked bar plots indicate the proportion of each CAR T-cell population at different time points, revealing an initial response phase involving CD8^+^ and double-negative CAR T-cells, followed by a long-term remission stage dominated by this CD4^+^ Ki-67^hi^ population. **d**, Ki67 expression in CD4^+^ CAR^+^ and CAR^-^ cells; and CD8^+^ CAR^+^ and CAR^-^ cells across time points. Statistical testing was performed using the Wilcoxon rank-sum test. Asterisks indicate significance levels. ****: p < 0.0001; ***: p < 0.001; **: p < 0.01; *: p < 0.05; ns = p > 0.05. **e**, Heatmap indicating z-score normalized expression of CyTOF markers across five major CAR T-cell clusters.

The CD4^+^ CAR T-cells were notable for a subpopulation highly expressing Ki-67, suggestive of a proliferative phenotype (**Fig. 2b**). Ki-67^hi^ CD4^+^ CAR T-cells steadily emerged as the dominant population in both patients: this population constituted 15.9% of CAR T-cells at month 1.8 in patient 1, increasing to 97.0% by year 9.3; and constituted 0.2% of CAR T-cells in patient 2 at month 2.4, increasing to 87.2% by year 7.2 (**Fig. 2b-c**). We assessed Ki-67 expression in the CD4^+^ CAR T-cells compared to the CAR^-^ T cells from these patients at matched time points, finding that this level of Ki-67 expression was strongly CAR T-cell specific (**Fig. 2d**). CD8^+^ CAR T-cells also exhibited a proliferative trend overall, but Ki-67 expression was generally lower and less robustly observed compared to the CD4^+^ CAR T-cell subset (**Fig. 2d**). These Ki-67^hi^ CD4^+^ T cells expressed a distinct marker profile, including activation markers CD38, HLA-DR, and CD95; transcription factors EOMES and TOX; checkpoint markers CTLA-4, LAG-3, TIGIT; and memory markers CD27 and CCR7 (**Fig. 2e**). Together, these data suggest two major phases of CAR T-cell therapy responses: an initial response phase dominated by cytotoxic CD8^+^ T cells and double-negative Helios^hi^ CAR T-cells, and a long-term remission phase dominated by a uniquely proliferative CD4^+^ CAR T-cell phenotype.

### Integrative single-cell analysis captures the landscape of RNA expression, protein expression, and TCR clonotype among CAR T-cells at 9.3 years post-infusion

Given the intriguing evidence of a distinct, pauci-clonal CD4^+^ CAR T-cell phenotype associated with long-term persistence, we sought to characterize the CAR T-cell clonality, protein expression, and whole-transcriptome profile of this population at a single-cell level. For this analysis, we selected patient 1 at year 9.3, the longest follow-up time point with sufficient cells available for this analysis. We generated joint single-cell TCR- and CITE-Seq libraries from CD3^+^ CAR^+^ DAPI^-^ sorted cells, obtaining a simultaneous readout of TCR clonotype, protein expression, and RNA expression for each cell. Despite the rarity of CAR T-cells at this time point, we obtained single-cell profiles from 1,437 T cells that passed our strict quality thresholds (**Fig. 3a**). By identifying single-cell RNA-Seq reads mapped to the 5’ CAR sequence, we identified 1,149 CAR T-cells, 288 of which were non-CAR expressing (“normal”) T cells that passed through our FACS gating (**Fig. 3b**).

**Fig. 3.**
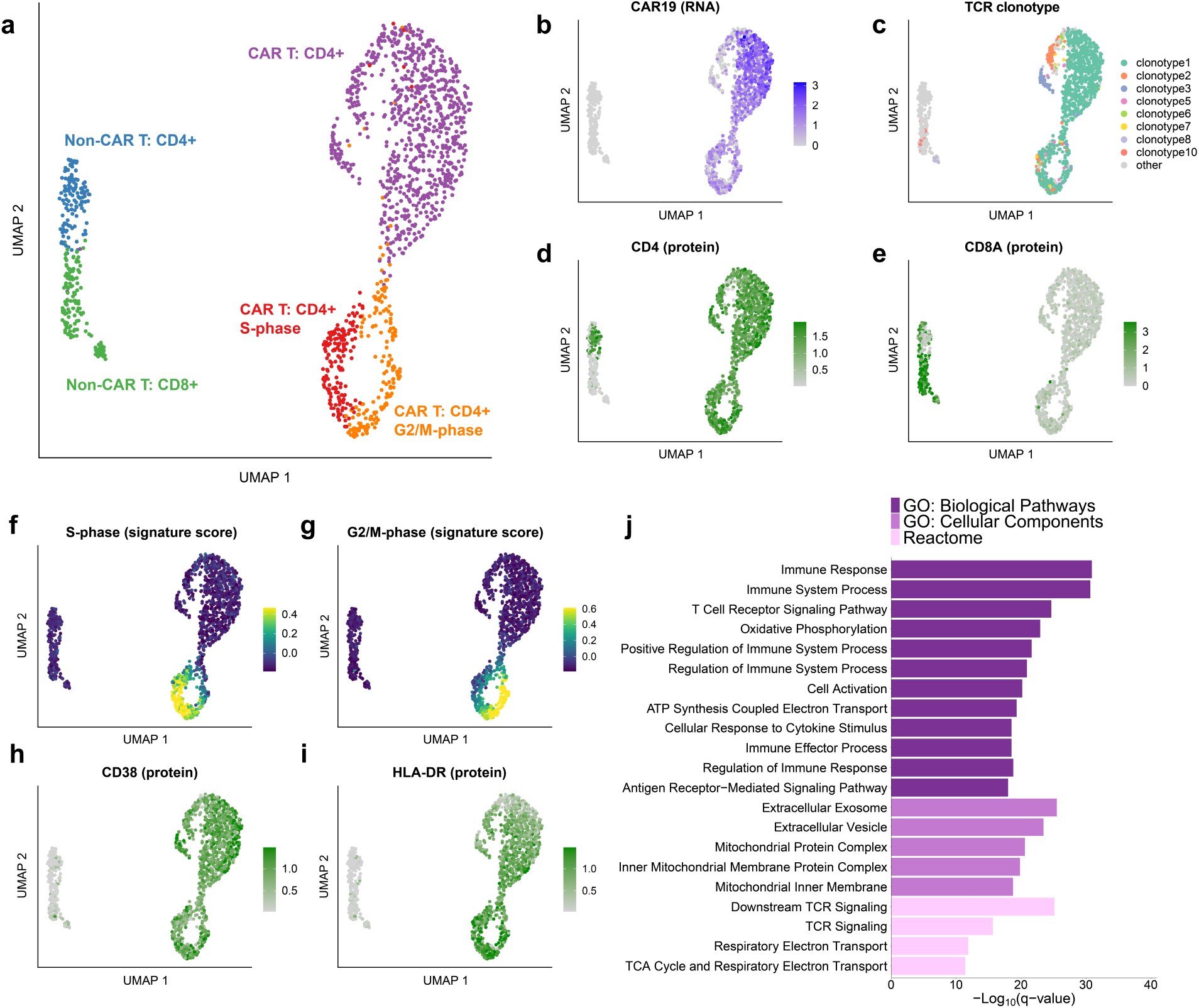
Integrative single-cell analysis reveals clonal expansion, proliferation, and activation in CAR T-cells from patient 1 at year 9.3. **a,** UMAP of 1,437 T cells sorted from peripheral blood from patient 1 at year 9. UMAP coordinates were computed using the single-cell RNA-Seq component of the 5’ TCR/CITE-Seq protocol. **b**, Normalized expression of the CD19BBz CAR construct detected from 5’ single-cell RNA-seq reads. **c**, UMAP colored by the eight detected TCR clonotypes with at least 10 cells each. Minor clusters with fewer than 10 detected cells were colored light gray. **d**, Normalized CITE-Seq antibody expression for the CD4 and (**e**) CD8A proteins. **f**, Cell cycle scores for cells in the S-phase or (**g**) G2/M phases. **h**, Normalized CITE-Seq antibody expression for the activation markers CD38 and (**i**) HLA-DR. **j**, Gene set enrichment analysis for genes significantly up-regulated in CD4^+^ CAR T-cells compared to CD4^+^ CAR^-^ T cells. Gene Ontology Biological Pathways and Cellular Components, and Reactome pathways were considered.

We reasoned that the non-CAR T-cells provide a valuable comparison to the CAR T-cell populations identified. Indeed, we found that CAR T-cells were markedly distinct from non-CAR T-cells with respect to clonal diversity, T cell subtype composition, metabolic and activation phenotype, and transcriptional regulation. While the normal T cells were clonally diverse (Shannon entropy = 6.88) with 170 unique clonotypes detected, the CAR T-cells demonstrated strong evidence of clonal dominance (Shannon entropy = 1.43), with the top 3 clonotypes constituting over 90% of CAR T-cells and only 27 unique clonotypes detected overall (**Fig. 3c**). Out of the 27 CAR T-cell clonotypes detected, 13 were detected at a previous time point from our prior TCR-Seq analysis from the first two years, including the most frequent clonotype seen at year 1.9 (indicated as clonotype 3 in **Fig. 3c**). Fourteen of the clonotypes, including the two most frequent clonotypes at year 9.3 (indicated as clonotypes 1 and 2 in **Fig. 3c**), were not observed at a previous time point (**Extended Data Fig. 4a**), consistent with our LVIS analysis that suggested the emergence of novel clones at the long-term follow-up time points. The normal T cells consisted of a relatively balanced mix of CD4^+^ and CD8^+^ T cell populations, whereas CD4^+^ CAR T-cells were sufficiently dominant at this time point that a CD8^+^ CAR T-cell cluster was not clearly identified (**Fig. 3d-e**).

CAR T-cells displayed evidence of ongoing proliferation at the transcriptomic level, supported by markers such as *PCNA*, *MCM6*, *TOP2A*, *CDK1*, *CCNB2*, and *MKI67* (**Extended Data Fig. 4b**) as well as from cell cycle scoring^13^ using gene sets to identify cells in S-phase or G2/M phase (**Fig. 3f-g**). Approximately 30% of CAR T-cells were observed to be in S, G2, or M-phase, compared to fewer than 7% of CAR non-expressing T cells (**Extended Data Fig. 4c-d**, Chi-squared p-value = 8.97e-15). This cell cycle effect was observed in CAR T-cells in a manner that was not clonotype-specific, with cell cycling seen across CAR T-cell clonotypes (**Extended Data Fig. 4e**).

### Functionally activated, cytotoxic characteristics among long-persisting CD4^+^ CAR T-cells

We next asked how CAR T-cells at this 9.3-year time point differed from their CAR^-^ counterparts with regard to cell surface phenotype and RNA expression. We were particularly curious whether these CD4^+^ CAR T-cells were cytotoxic in nature, or if they exhibited traits of T cell exhaustion. CITE-Seq antibody expression levels suggested that the CAR T-cells exhibited a distinct cell surface phenotype compared to CAR^-^ T cells, including expression of activation markers CD38 and HLA-DR (**Fig. 3h-i**). The CAR T-cells also expressed the inhibitory receptors PD-1, TIM-3, LAG-3, and TIGIT, which have been associated with both activated and exhausted T cell states (**Extended Data Fig. 5a-b**). We identified 645 differentially expressed genes in the comparison between CAR T-cells in G1 phase compared to the normal CD4^+^ T cell population in G1 phase (**Extended Data Fig. 4f**, **Supplementary Table 1**, Bonferroni adjusted p-value < 0.001), with up-regulated genes enriched for an effector CD4^+^ T cell signature (FDR = 0.024, **Extended Data Fig. 4g**); in contrast, only thirty-three genes were differentially expressed between the top three clonotypes (**Extended Data Fig. 4h**). We observed significant enrichment in T cell activation, TCR signaling, oxidative phosphorylation, vesicle components, and mitochondrial protein complexes in the CAR T-cells (**Fig. 3j**). Consistent with a functionally activated rather than exhausted phenotype, we observed upregulation of genes encoding for cytokines IL-10 and IL-32 among CAR T-cells (**Fig. 4a**); this functional activity appeared to be associated with antigen-driven signaling through the CAR, as TCR signaling and antigen-mediated signaling were the top-enriched signaling pathways (**Fig. 3j; 4b**). Intriguingly, we observed a distinct expression profile of cytotoxic enzymes in the CD4^+^ CAR T-cells. *GZMK* and *GZMA* were among the top four up-regulated genes among the CD4^+^ CAR T-cells (**Fig. 4c; Extended Data Fig. 4f**), suggestive of direct cytotoxic function. Perforin gene *PRF1* was additionally up-regulated, along with enrichment of vesicle cellular components that may be involved in cytotoxic granules (**Fig. 3j; Fig. 4c**); whereas *GZMB* and *GZMH* were not highly expressed.

**Fig. 4.**
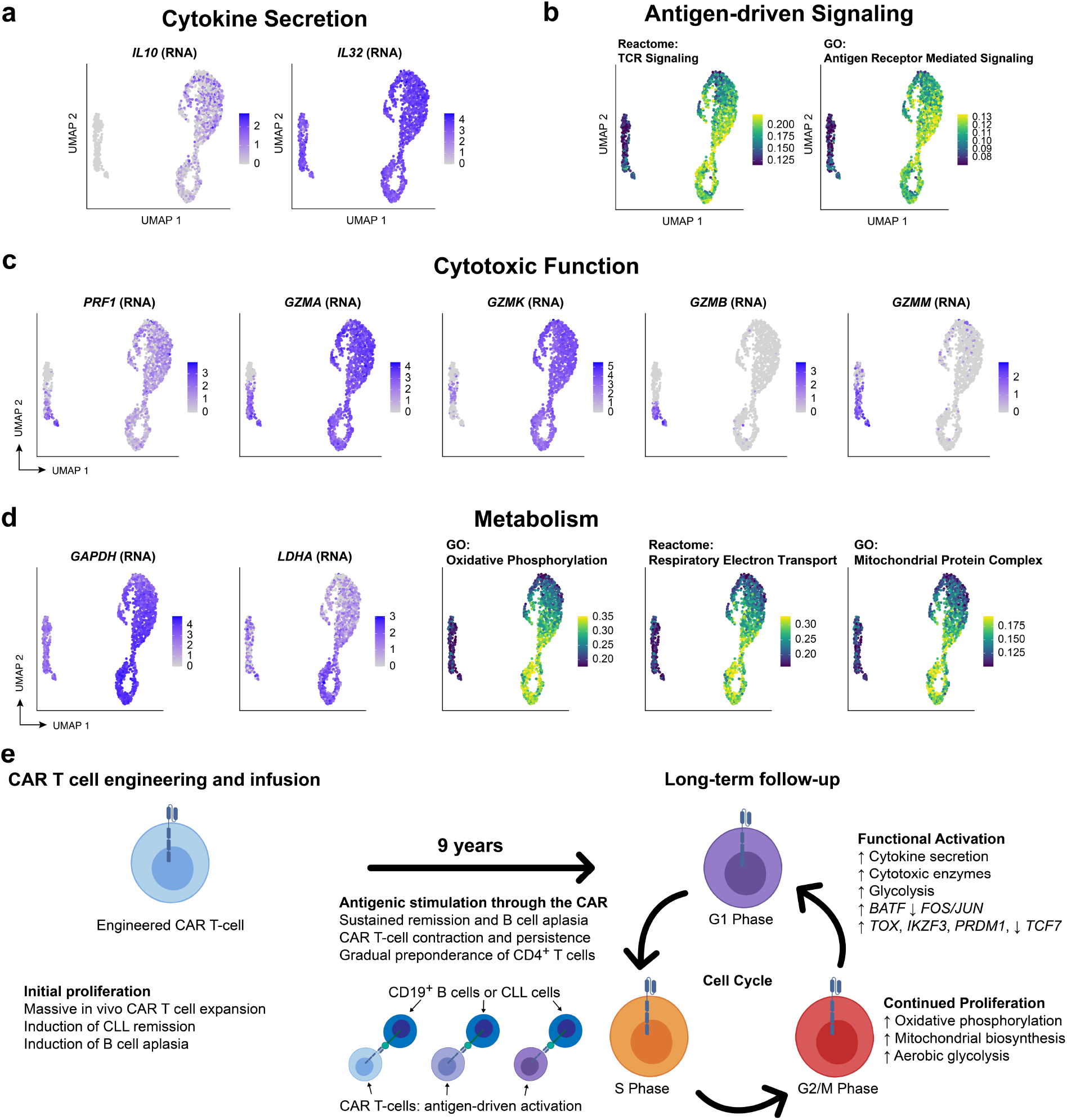
Evidence of functional activation, metabolic reprogramming, and antigen-driven signaling among CAR T-cells from patient 1 at year 9. **a**, UMAP indicating upregulated RNA expression of cytokine genes *IL1*0 and *IL32* among CAR T-cells compared to normal T cells. **b**, AUCell scores for Reactome TCR signaling and Gene Ontology Antigen Receptor Mediated Signaling pathways. **c**, UMAP indicating upregulated RNA expression of cytotoxic genes *PRF1*, *GZMA*, and *GZMK* genes among CD4^+^ T cells, with *GZMB* and *GZMM* expressed only in the normal CD8^+^ T cells. **d**, RNA expression of key glycolytic gene *GAPDH* and fermentative glycolysis gene *LDHA*; AUCell scores for oxidative phosphorylation, respiratory electron transport, and mitochondrial protein complex up-regulated among CAR T-cells, particularly in the active cell cycling phases. **e**, Proposed model of mechanistic basis of sustained remission with B cell aplasia mediated by few, metabolically active but immune checkpoint inhibitor-restrained CAR T-cell clones.

The CAR T-cells exhibited a metabolic phenotype consistent with increased glycolysis in all cell cycle phases, and oxidative phosphorylation and aerobic glycolysis in the cell cycling states (**Fig. 4d**). Glycolytic gene *GAPDH* was upregulated in all cell cycle stages of the CAR T-cells compared to CAR^-^ T cells, and the gene encoding for lactate dehydrogenase A (*LDHA*) was up-regulated among the actively cycling cells (**Fig. 4d-e**), suggestive of Warburg-like aerobic glycolysis in the CAR T-cells. Single-cell gene signature scoring with AUCell^14^ revealed that oxidative phosphorylation and respiratory electron transport pathways were upregulated particularly in the actively cycling S/G2/M phase cells, as well as mitochondrial protein cellular components (**Fig. 4d**), suggesting that mitochondria-dependent oxidative phosphorylation and aerobic glycolysis provide critical metabolic support for the proliferation of the CAR T-cells.

Finally, we sought to characterize the transcriptional regulation underlying this functionally active CAR T-cell phenotype in patient 1 at the 9.3-year time point. Differential expression analysis revealed 18 significantly differentially expressed transcription factors (**Extended Data Fig. 6a**, Bonferroni adjusted p-value < 0.001), and single-cell regulon scores generated using GENIE3^15^ and AUCell showed strong correlation between transcription factor regulons (**Extended Data Fig. 6b**). *TCF7* expression was downregulated among the CAR T-cells compared to the non-activated CAR^-^ cells, whereas the transcription factors *TOX*, *IKZF3*, *EOMES*, and *PRDM1* were associated with the activated CD4^+^ CAR T-cell state (**Extended Data Fig. 6c**). We observed significant differential expression of multiple AP-1 transcription factors, including downregulation of *FOS*, *JUNB*, and *JUN* expression and upregulation of *BATF* (**Extended Data Fig. 6d**).

## Discussion

Little is known about the fate of long-term persisting CAR T-cells in patients with durable remissions. Here, we report the functional and molecular characterization of the longest persisting anti-CD19 CAR T-cells reported to date. We observed two distinct phases of the anti-leukemia response, beginning with an initial response phase characterized by cytotoxic CD8^+^ or double-negative Helios^hi^ CAR T-cells, followed by the predominance of a proliferative CD4^+^ CAR T-cell population in both patients in the ensuing years. The observed CD4^-^CD8^-^ Helios^hi^ CAR T population resembled recently reported Helios-expressing double-negative T cells derived from CD8^+^ T cells^16^, with NKT or gamma-delta T cells as other possible identities. CITE-Seq analysis demonstrated that the long-persisting CD4^+^ CAR T-cells exhibited evidence of ongoing proliferation, cytokine expression, and metabolic activity that strongly suggested that they remained functionally active rather than exhausted. The CAR T-cells detected in patient 1 at year 9 were exclusively CD4^+^, a surprising finding that led us to rethink the possibility that CD4^+^ T cells, not CD8^+^ T cells, may be primarily responsible for cytotoxicity against CD19-expressing cells at these time points. Strongly up-regulated antigen-mediated signaling pathways and up-regulation of *GZMK*, *GZMA*, and *PRF1* supported this notion, as well as a similarity to a recently reported cytotoxic CD4_GZMK_ T cells population in a study of bladder cancer^17^. These findings offer intriguing new insights into the nature of long-term CAR T-cell signaling and persistence among these unique patients.

## Materials and Methods

### Study Design

Both patients were enrolled on the trial designed to determine safety and feasibility of anti-CD19 CAR T-cell manufacturing and infusion in patients with relapsed or refractory B-cell malignancies^18^. The trial (ClinicalTrials.gov number, NCT01029366) was approved by the institutional review board at the University of Pennsylvania and conducted in accordance with the protocol^3^. Autologous T cells were collected by apheresis, enriched and activated using anti-CD3 and anti-CD28-coated polystyrene beads (Dynal) and transduced with the murine, FMC63-based chimeric antigen receptor carrying 4-1BB and CD3-zeta signaling domains as previously reported^18^. Patients 1 and 2 were infused with a total dose of 1.1 x 10^9^ (1.6 x 10^7^/kg) and 1.4 x 10^7^ (1.46×10^5^/kg) CAR T-cells, respectively, on three consecutive days and monitored for CAR T-cell engraftment, tumor dynamics, and systemic inflammatory responses using previously reported qualified flow cytometry, quantitative PCR, and Luminex assays^3,8^. All ethical guidelines were followed. No commercial sponsor was involved in the study.

### Correlative studies

Patient samples were acquired prior to manufacturing, from the infusion product itself, and after infusion for the studies described here. Our routine pipeline of correlative assays includes a) a flow cytometry-based assay using a non-commercially available, in-house generated anti-CAR19 idiotype monoclonal antibody (mAb) conjugated to either Alexa Fluor 647 (kind gift from Dr. Laurence Cooper, MD Anderson, Houston, TX) or PE (Novartis Institute for Biomedical Research); b) a quantitative PCR assay with primers spanning the 4-1BB and CD3ζ chimeric molecule; and c) a flow-cytometry-based assay to quantify B cells and leukemic cells. All assays have been qualified prior to implementation in the clinical monitoring and extensively used^8,19^. All routine correlative assays were qualified prior to implementation, and carried out at time points defined by the clinical protocol in parallel with disease response evaluations. In addition, leukemia response to CTL019 in these patients was assessed via deep sequencing of immunoglobulin heavy chain rearrangements as previously described^3,12^.

### Cytometry by Time-of-Flight (CyTOF)

Antibodies for mass cytometry were obtained as pre-conjugated metal-tagged antibodies from Fluidigm or generated in-house by conjugating unlabeled purified antibodies to isotope-loaded polymers using MAXPAR kits (Fluidigm). Antibodies generated in-house were diluted in antibody stabilization buffer (Candor Bioscience). In total, 35 mAb were used to identify T cell expressed surface and intracellular proteins in conjunction with five mAb and dead cell exclusion dye to remove non-T cells and artifacts from analysis.

Cryopreserved PBMCs were thawed with thawing medium containing 10% of FBS in RPMI supplemented with benzonase (0.5U/ml). Thawed cells were washed with PBS once and transferred to a 96-well U-bottom tissue culture plate. Single cell suspensions were pelleted, and incubated with the Cisplatin solution (Final concentration: 25uM) for 1min at room temperature (rT) for live/dead discrimination. The reactions were quenched with PBS/1% FBS (flow buffer) and centrifuged for 4 min at 500x g. Cell pellets were resuspended in surface antibody cocktail (adjusted to 50ul final volume flow buffer, incubated for 20min at 4°C, and washed twice in flow buffer. The cells were fixed and permeabilized in 50ul of FOXP3 Fixation/Permeabilization working solution (FOXP3 staining buffer set, eBioscience), and stained intracellularly for 30 min at 4°C. The cells were further washed twice with 1x permeabilization buffer (eBioscience) before fixation in 1.6% PFA solution (Electron Microscopy Sciences) containing 125 nM Iridium (Fluidigm) overnight at 4°C. Prior to data acquisition on the Helios mass cytometer (Fluidigm), cells were washed twice in PBS (without Ca^2+^ and Mg^2+^) and once in deionized H_2_O. Immediately prior to sample acquisition, cells were resuspended with deionized H_2_O containing the bead standard at a concentration 1-2 × 10^4^ beads per mL. Samples were acquired using a bead-based normalization of CyTOF data by using Nolan lab normalizer available through https://github.com/nolanlab/bead-normalization/releases.

### T cell Receptor Vβ (TRB) deep sequencing

CART19 cells were purified from early (first 24 months) post-infusion specimens for TRB deep sequencing (TCR-seq) using a Becton Dickinson Aria II flow cytometer. Genomic DNA was isolated using the DNeasy Blood and Tissue Kit (Qiagen) and TCR-seq was carried out by (Adaptive Biotechnologies). Only productive TCR rearrangements were used in the assessment of TCR clonotype frequencies.

### Lentiviral vector integration site analysis

Vector integration sites were detected from genomic DNA as described previously^11,12^. Genomic sequences were aligned to the human genome by BLAT (hg38, version 35, >95% identity) and statistical methods for analyzing integration site distributions were carried out as previously described^20^. The SonicAbundance method was used to infer the abundance of cell clones from integration site data^21^, and annotatr^22^ was used to annotate the genomic location of the integration sites. All samples were analyzed independently in quadruplicate to suppress founder effects in the PCR and stochastics of sampling.

### CyTOF data processing

Cytobank^23^ was utilized to computationally gate live CAR^+^ CD3^+^ T cells. Gating was performed as previously described^24^ to isolate live singlets, which were further processed by removing CD14/CD19 positive cells, selecting CD45^+^/CD3^+^ cells, and gating CAR^+^ and CAR^-^ using healthy-donor derived PBMCs to set the CAR^+^ gate. Pre-processed data were imported into Seurat. Expression values are arcsinh transformed with a cofactor of 5, and matrices were concatenated. UMAP projections were computed using the top 10 PCs and 20 nearest neighbors. Initial clustering was performed with the Louvain algorithm implemented in Seurat with a resolution parameter of 0.8, and 14 clusters were merged to five meta-clusters based on similar marker expression in Extended Data Fig. 3b. In order to avoid the impact of outliers on color scales, color scales for UMAPs were defined from a range of zero to the 99^th^ percentile of expression. Antibody clones are provided in **Supplementary Table 2**.

### Single-cell immune, proteome, and transcriptome profiling by 5’ CITE-Seq and TRB V(D)J sequencing

Thawed patient PBMCs were washed with 1x PBS (Life Technologies) and resuspended with antibody staining buffer containing human TruStain FcX (Biolegend). Cells were incubated with TotalSeq-C, anti-CD3 (BioLegend) and CAR (custom generated, Novartis) antibodies for 30 min at 4°C. Stained cells were washed with antibody staining buffer three times and filtered through Flowmi. Before sorting, DAPI (Thermal Fisher) was added to sort single CD3^+^ CAR^+^ nuclei. Antibody clones are provided in **Supplementary Table 2**.

Sorted single nuclei were loaded on a Chromium Chip G (10x Genomics) according to the manufacturer’s instructions for processing with the Chromium Next GEM Single Cell 5’ Library & Gel Bead Kit v1.1. TCR single-cell library and cell surface protein libraries were subsequently prepared from the same cells with the Chromium Single Cell V(D)J Enrichment Kit, Human T Cell and 5’ Feature Barcode Library Kit, separately. TCR single-cell library, 5’ gene Expression library and cell surface protein library were pooled with a molar ratio 1:10:1 for sequencing on Illumina NextSeq 550 with 26×91 bp, aiming for 50,000 read pairs per cell for 5’ gene expression library and 5,000 read pairs per cell for both TCR single-cell library and cell surface protein library.

### Analysis of CITE-Seq and single-cell V(D)J sequencing data

Demultiplexing and alignment of CITE-Seq RNA and antibody-derived tag sequences were performed with cellranger v3.1.0 using the Gencode v32 reference modified to contain the 5’ CAR sequence. TCR sequences were identified with cellranger vdj, and clonotypes were defined based on TRB CDR sequences. Low-quality cells were computationally filtered by retaining only those cells with between 200 and 5000 genes in the scRNA-Seq data, less than 5% mitochondrial RNA, and containing a detectable TCR sequence. Centered log-ratio normalization was performed on CITE-Seq antibody-derived tag counts. Single-cell RNA-Seq count matrices were log-normalized, and the top 2000 variable genes were identified with the variance-stabilizing transformation with Seurat v3.2.0^25^. Dimensionality reduction was performed using UMAP on the top 15 principal components using 30 neighbors and 2 components.

Differential expression analysis on single-cell data was performed using the Wilcoxon rank sum test with Bonferroni multiple testing correction. Pathway enrichment analysis on differentially expressed genes was performed using gprofiler2^26^, and single-cell pathway enrichment scores were defined using AUCell v1.6.1^14^. For visualization of CITE-Seq protein markers and AUCell enrichment scores, maximum values of color gradients were defined using the 5^th^ and 95^th^ percentile values in order to reduce the impact of outlier values. GENIE3^15^ was used to define a transcriptional regulatory network, and regulons of transcription factors were defined by identifying target genes with an edge score greater than 0.005 and a positive expression correlation.

## Acknowledgments

The authors acknowledge the contributions of the Translational and Correlative Studies Laboratory for providing standardized flow cytometric, qPCR, and cytokine multiplexing analyses, and biobanking of patient specimens on CAR T-cell trials; the clinical Cell and Vaccine Production Facility for cell processing and biobanking; the Human Immunology Core for providing healthy donor cells and analytical support; the Flow Cytometry Core for flow cytometry equipment maintenance; and the Stem Cell & Xenograft Core for providing patient specimens.

## Data Availability Statement

Raw sequencing data for this study are in preparation for submission to dbGaP (accession number pending).

## Author contributions

Author contributions: JJM, GMC, MW, DLP, PG, SB, IPM, CLN, SM, LT, IK, MG, DEA, FDB, SFL, KT, and CHJ performed experiments or analyzed the data. All authors helped design the experiments and contribute to data interpretation. JJM, GMC, MW, DLP, KT, and CHJ wrote the manuscript.

## Extended Data Figures and Tables

**Extended Data Table 1.**
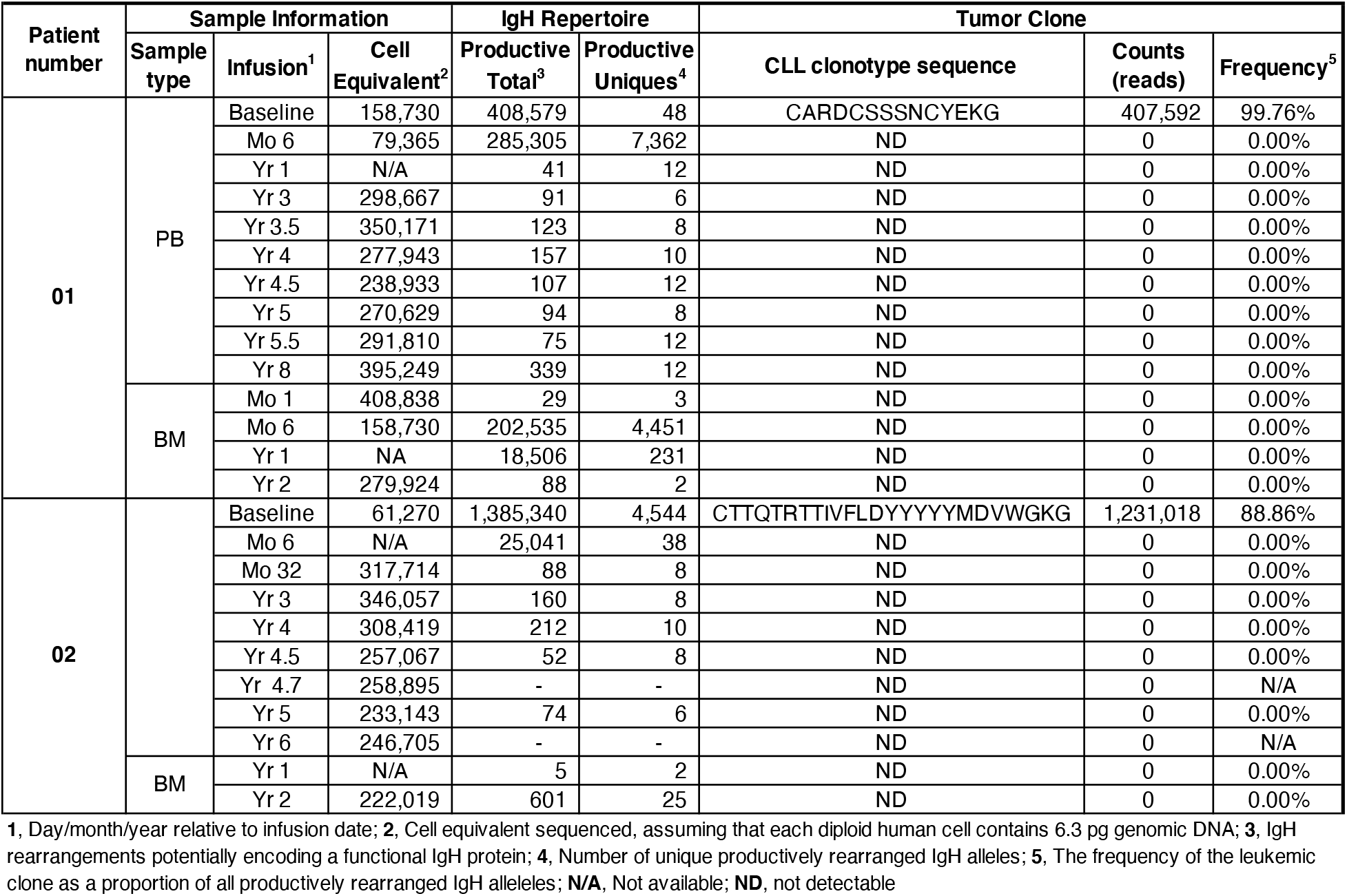
Immunoglobulin heavy chain rearrangement deep sequencing shows persistent deep molecular remission for both patients.

**Extended Data Fig. 1.**
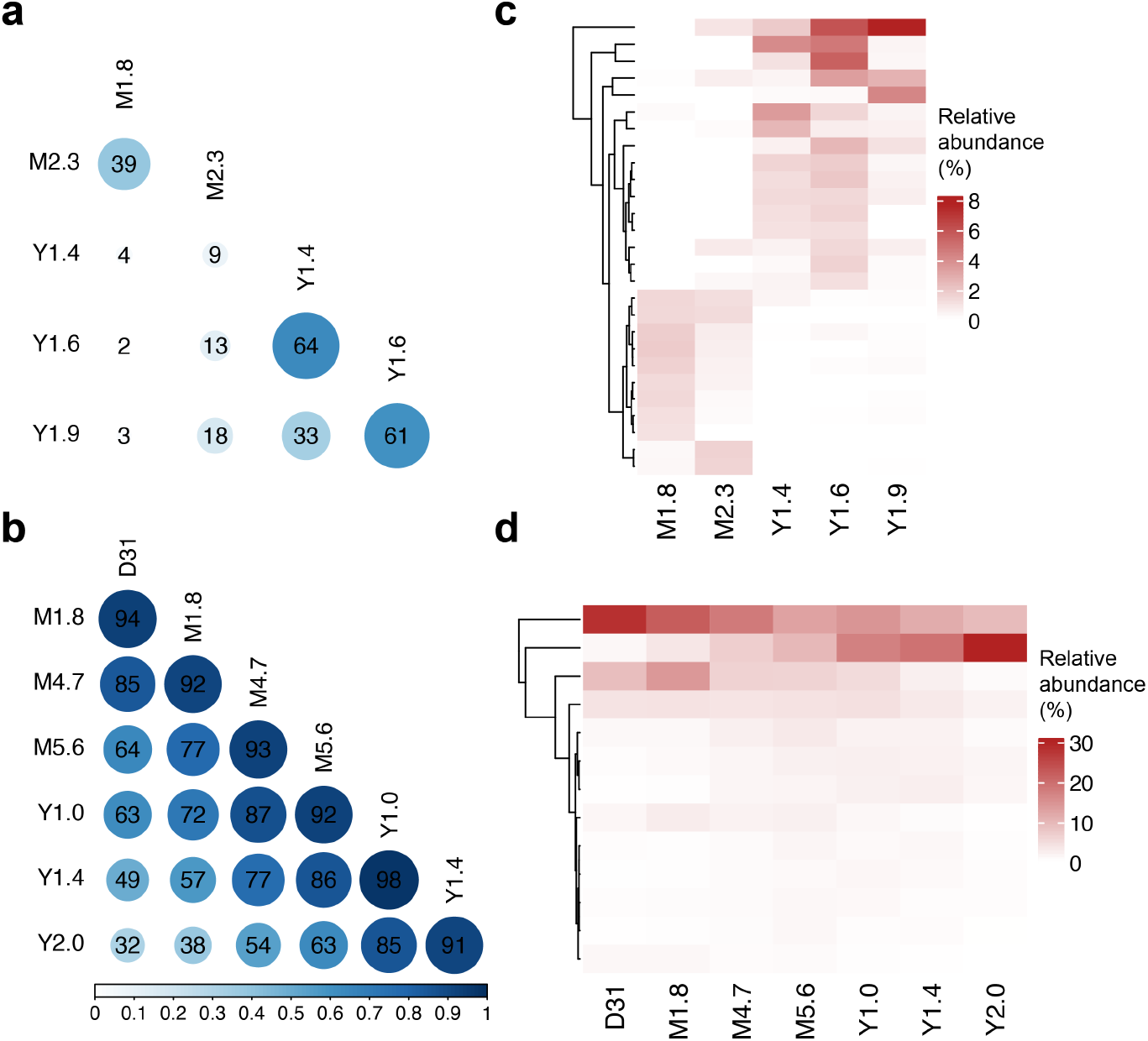
Clonal evolution for patient 1 and 2 based on TCR sequencing data. Pairwise Morisita’s overlap index was computed between all timepoints (row and column labels) for patient 1 (**a**) and 2 (**b**). TCR clones (rows) with maximum abundance > 1% across time points were retained and tracked over time for patient 1 (**c**) and 2 (**d**).

**Extended Data Fig. 2.**
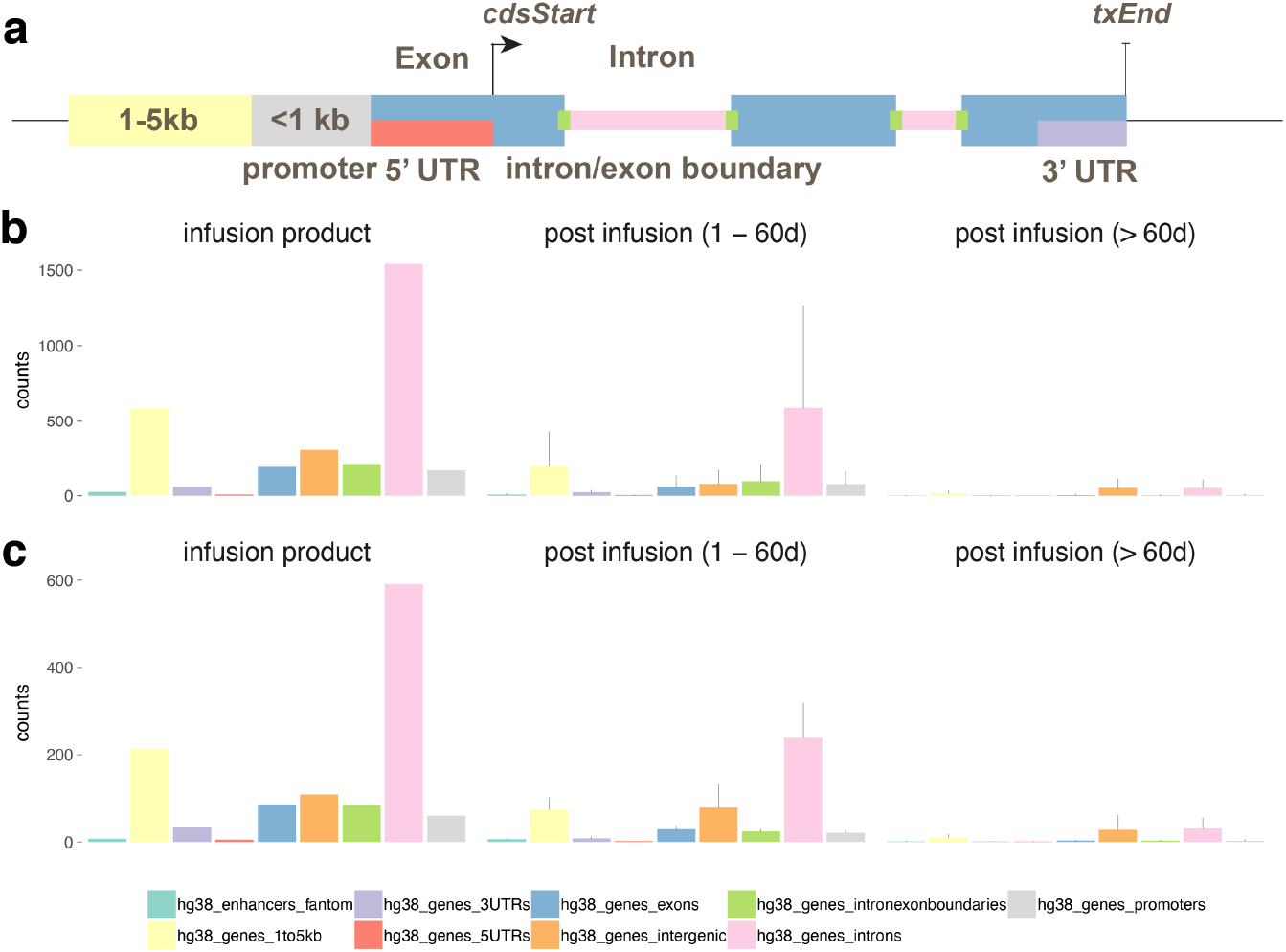
Genomic annotation of integration sites for infusion product, post infusion timepoints < 60d and > 60d. **a**, graphic of the annotation scheme. The integration sites from patient 1 (**b**) and 2 (**c**) were annotated based on its position relative to known genes (UCSC hg38) and permissive enhancers (FANTOM 5). The counts of integration sites that fall into each annotation category in infusion product were summarized (left). The mean and standard deviation of the number of sites for each category were also computed for 1 - 60 days post infusion (middle) and > 60 days post infusion (right).

**Extended Data Fig. 3.**
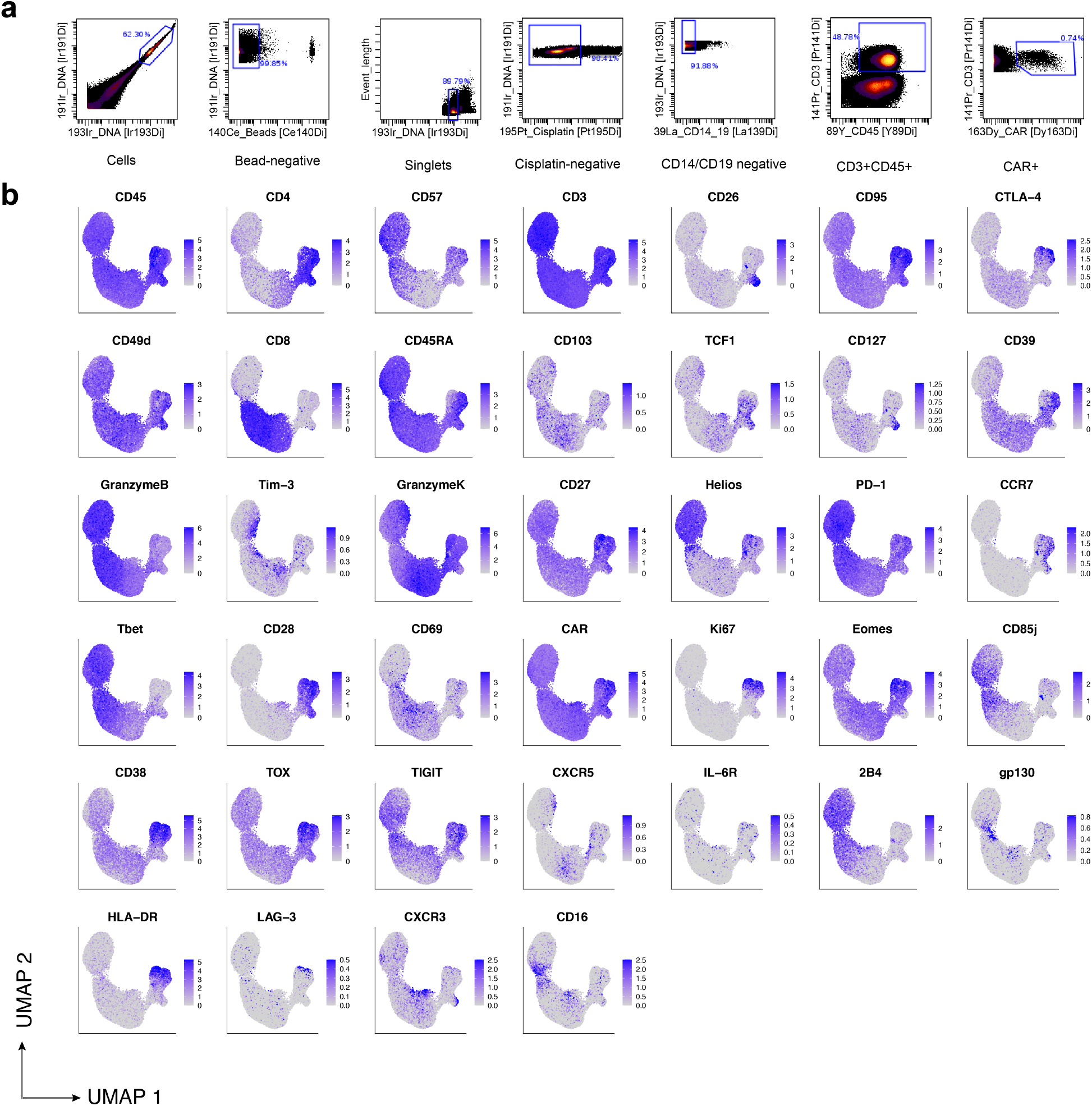
Gating strategy and CyTOF marker expression profiles. **a**, Gating strategy performed computationally on CyTOF data to filter to CD3^+^CAR^+^ T cells for downstream analysis. **b**, Protein expression of our CyTOF panel depicted on a single-cell basis on our UMAP.

**Extended Data Fig. 4.**
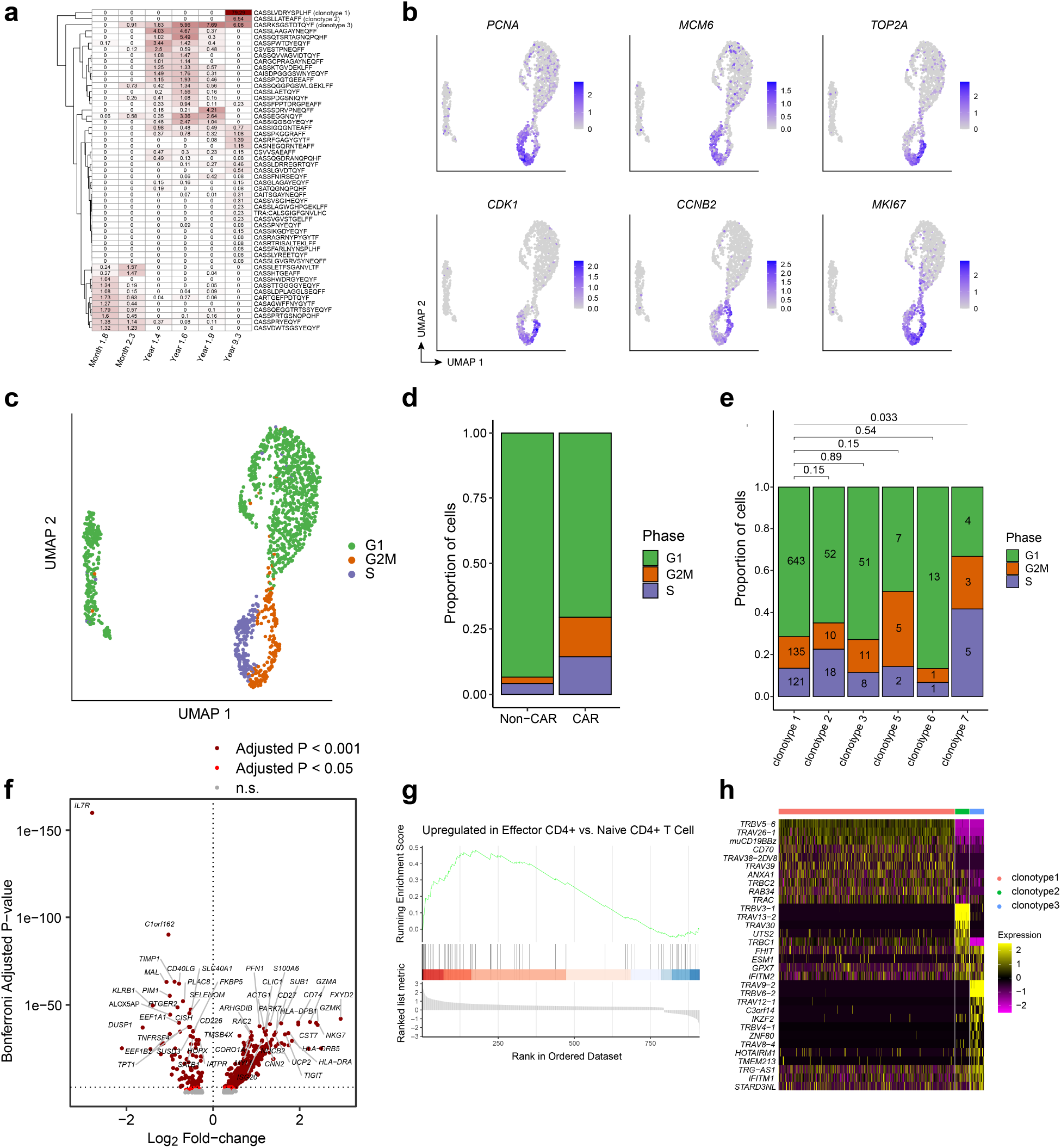
Analysis of CAR T-cell clonotype, cell cycle, and differential gene expression from patient 1 at year 9. **a**, Heatmap showing the relative frequencies of TCR clonotypes at the 2-month, 3-month, 15-month, 18-month, 21-month, and 9-year time points. Note that the first five columns were estimated from bulk TCR sequencing, whereas the rightmost column was estimated from the single-cell TCR/CITE-Seq data from year 9. **b**, UMAPs indicating strong up-regulation of RNA expression of cell cycle genes. **c**, UMAP colored by cell cycle phase using Seurat. **d**, Proportions of cells in each cell cycle phase, compared between CAR^-^ T and CAR^+^ T cells. Chi-squared p-value = 8.97e-15. **e**, Proportions of cells in each cell cycle phase, compared between the top six CAR T-cell clonotypes. Pairwise statistical significance was assessed with the Chi-Squared test, and multiple-testing correction was performed using the Benjamini-Hochberg method. Numbers within the bars indicate the number of cells observed. **f**, Volcano plot indicating genes up-regulated in CAR T-cells compared to normal CD4^+^ T cells (rightward direction) and genes down-regulated in CAR T-cells compared to normal CD4^+^ T cells (leftward direction). Differentially expressed genes were determined using the Wilcoxon rank-sum test with a Bonferroni-adjusted p-value cutoff of 0.001 (dark red) and 0.05 (red). **g**, Gene Set Enrichment Analysis plot for the effector CD4^+^ gene signature. **h**, Heatmap indicating normalized gene expression values for the 32 differentially expressed genes with a Bonferroni-adjusted p-value cutoff of 0.001.

**Extended Data Fig. 5.**
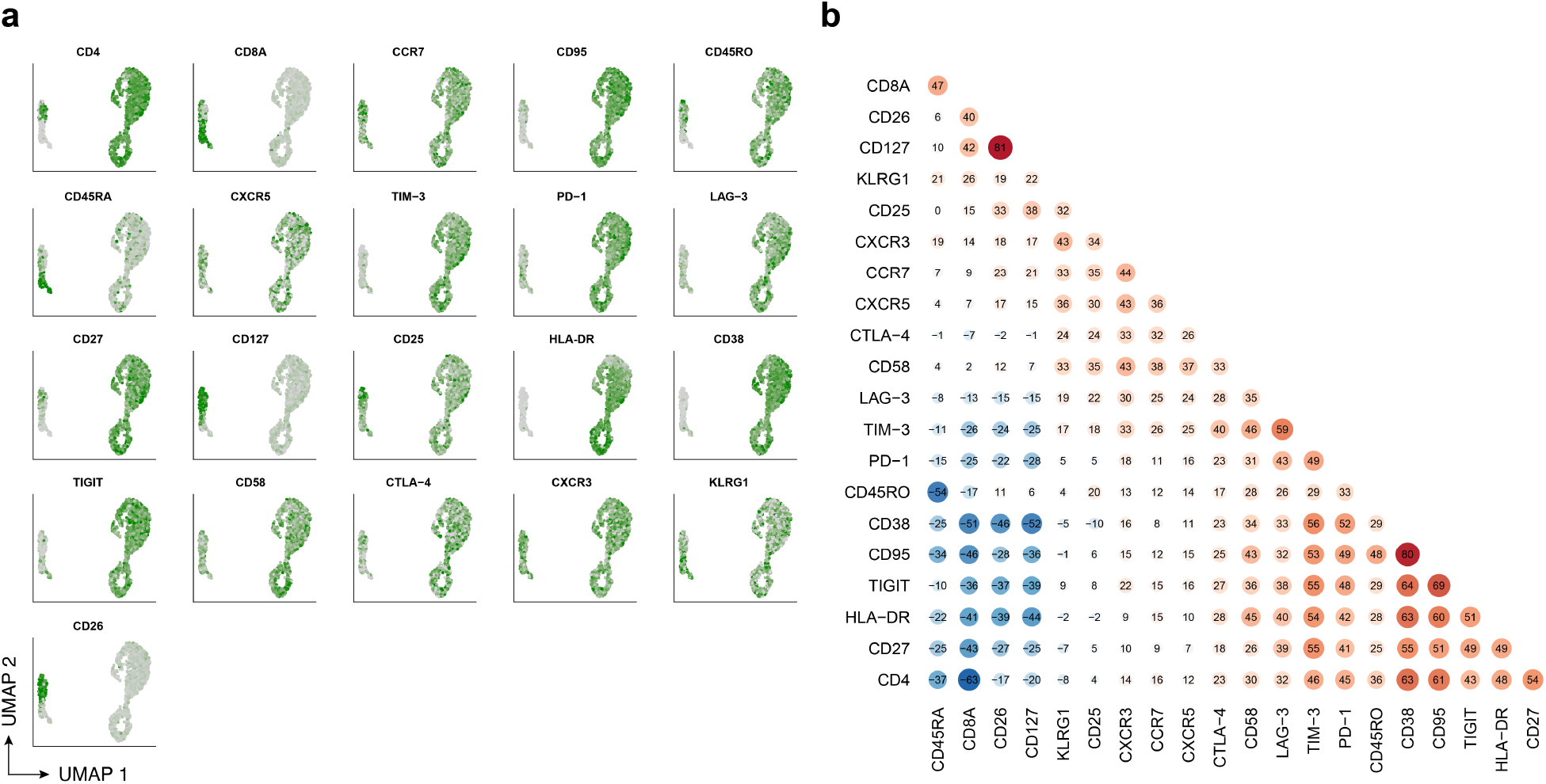
CITE-Seq protein expression and correlation for patient 1 at year 9. **a**, UMAP colored by normalized expression of CITE-Seq protein expression determined by antibody-derived tags. **b**, Pairwise Spearman correlations between CITE-Seq protein expression values across cells.

**Extended Data Fig. 6.**
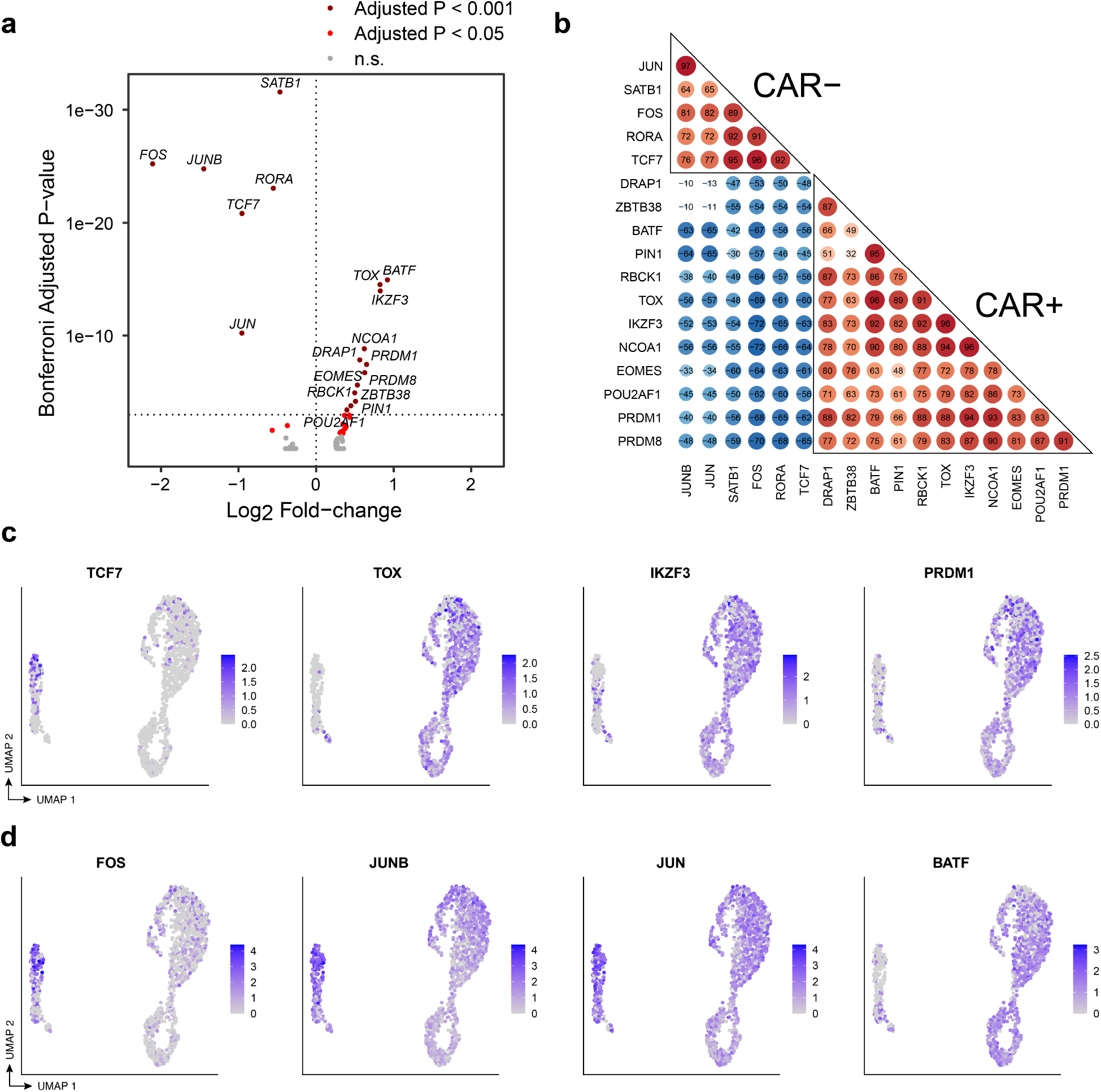
Transcriptional regulation of CAR T-cells in patient 1 at year 9. **a**, Volcano plot indicating transcription factors (TFs) up-regulated in CAR T-cells compared to normal CD4^+^ T cells (rightward direction) and TFs down-regulated in CAR T-cells compared to normal CD4^+^ T cells (leftward direction). Differentially expressed TFs were determined using the Wilcoxon rank-sum test with a Bonferroni-adjusted p-value cutoff of 0.001 (dark red) and 0.05 (red). **b**, Pairwise correlation of TF regulon scores determined by GENIE3 and AUCell in the comparison between CAR T-cells and CD4^+^ CAR^-^ T cells. **c**, UMAP indicating RNA expression of selected differentially expressed TFs *TCF7*, *TOX*, *IKZF3*, and *PRDM1*. **d**, UMAP indicating RNA expression of differentially expressed AP-1 TFs, *FOS*, *JUNB*, *JUN*, and *BATF*.

